# Central and peripheral tau retention modulated by an anti-tau antibody

**DOI:** 10.1101/2023.08.17.553682

**Authors:** Alexander Solorzano, Molly Brady, Nemil Bhatt, Angelique Johnson, Brooke Burgess, Hannah Leyva, Nicha Puangmalai, Cynthia Jerez, Ronald Wood, Rakez Kayed, Rashid Deane

## Abstract

Tau protein blood levels dependent on its distribution to peripheral organs and possible elimination from the body. Thus, the peripheral distribution of CSF-derived tau protein was explored, especially since there is a transition to blood-based biomarkers and the emerging idea that tau pathology may spread beyond brain. Near infrared fluorescence (NIRF) was mainly used to analyze tau (tau-NIRF) distribution after its intracisternal or intravenous injection. There was a striking uptake of blood- or CSF-derived tau-NIRF protein by the skeletal structures, liver, small intestine (duodenum), gall bladder, kidneys, urinary bladder, lymph nodes, heart, and spleen. In aging and in older APP/PS1 mice, tau uptake in regions, such as the brain, liver, and skeleton, was increased. In bone (femur) injected tau protein was associated with integrin-binding sialoprotein (IBSP), a major non-collagenous glycoprotein that is associated with mineralization. Tau-NIRF was cleared slowly from CSF via mainly across the cribriform plate, and cervical lymph nodes. In brain, some of the CSF injected tau protein was associated with NeuN-positive and PDGFRý-positive cells, which may explain its retention. The presence of tau in the bladders suggested excretion routes of tau. CSF anti-tau antibody increased CSF tau clearance, while blood anti-tau antibody decreased tau accumulation in the femur but not in liver, kidney, and spleen. Thus, the data show a body-wide distribution and retention of CSF-derived tau protein, which increased with aging and in older APP/PS1 mice. Further work is needed to elucidate the relevance of tau accumulation in each organ to tauopathy.

## Introduction

Tau protein, an intracellular microtubule-associated protein, is a soluble protein that promotes tubulin assembly into microtubules and its stability^1–5^. Tau is abundantly expressed in neurons where it is involved in axonal transport, and as a major component of the cytoskeleton provides structural support to neuron^3,6,7^. In pathological conditions, such as Alzheimer’s disease (AD), abnormal accumulation of tau protein is associated with the formation of neurofibrillary tangles (NFTs), and microtubules disruption in neurons^1,2,5,8–10^. In normal brains, wild type tau protein is released into the synaptic cleft and interstitial fluid (ISF) during neuronal activity^11–16^, independent of cell death^1–6,17,18^. Tau can then distribute to surrounding cells, including post-synaptic neurons, which is associated with the synaptic transmission of tau^19–23^. Tau is also distributed to the vasculature, and via the bulk flow of ISF to CSF, which drains into blood at various cerebrospinal fluid (CSF) outflow sites along the CSF bulk flow pathways^24^.

There are several reports on the correlation between brain tau levels with that of CSF and blood serum and/or plasma^25–32^. Tau in blood is likely to be distributed to peripheral tissues and eliminate via the kidneys. Indeed, earlier reports have alluded to this by using immunohistochemistry (ICH), Western blot (WB) analysis or ELISA^33–40^. In AD human peripheral tissues, tau levels in the submandibular gland were greater than that of sigmoid colon, followed by liver, scalp and abdominal skin^40^. However, tau is widely expressed in tissues^41–43^. Thus, methods, such as IHC, WB and ELISA, may not distinguish between peripheral tissue-derived tau proteins and CNS-derived tau proteins. There is a need to further explore the CNS and peripheral distribution of CSF-derived tau, especially since there is a transition to blood-based biomarkers and the emerging idea that tau pathology may spread beyond brain.

To address this critical gap in knowledge, we mainly used monomeric tau protein conjugated to a near infrared fluorescence (NIRF) molecule, which provides a better contrast between the target and the natural background, as we have reported^24,44^. Tau conjugated to Alexa-555 was used to map its distribution and colocalization with tissues^24,44^. Our data show that tau was retained both with the CNS and peripheral organs, and this was greater with aging and in older APP/PS1 mice, an AD mouse model. Surprisingly, there was a striking body-wide distribution of blood- and CSF-derived tau-NIRF, including the skeletal structure, liver, small intestine (duodenum), kidneys, brain, lymph nodes, heart, and spleen. In aging and in APP/PS1 mice, tau was retained in brain, femur, and liver. In CSF, an anti-tau antibody increased tau clearance from CSF. However, in blood it differentially affected tau distribution to peripheral tissues by reducing its accumulation in the femur, but not affecting uptake in liver, kidney, and spleen. Further work is needed on the relevance of tau retention in organs, which may provide a better understanding of degenerative diseases/tauopathies, and whether this contributes to the concept for age-related degeneration.

## Results

### Multiorgan distribution CSF-derived tau protein includes the skeleton

Recently, we reported that the main routes for the CSF outflow were via the olfactory/nasal (across the cribriform plate) and spinal pathways using real-time in vivo kinetic analysis of bovine serum albumin (BSA-NIRF) to visualize CSF elimination ^24^. Tau-NIRF was intracisternally infused into mice and its signal intensity recorded continuously every 4 sec for up to 2.5 hrs from the nose to the end of the spine with the mice in a prone position (**Fig. 1a**). The main CSF outflow sites from the cisterna magna were the nasal/olfactory bulb (NOB) and spinal column (SP). For these regions, after a delay, there were a rapid increase in tau-NIRF signal to a deflection point at about 30 mins followed by a slower progressive linear increase (**Fig. 1b**). This pattern of intensity profile curve was unlike our report for BSA, which peaked after about 30 mins followed by a disappearance phase, which represents the elimination phase ^24^. The data suggested that there was likely retention of tau but not BSA. At the NOB, the area under the curve (AUC) was significant greater in the young mice (Y) compared to that of the older (O) mice (**Fig.1c**). Likewise, the AUC was significantly greater for the older APP/PS1 (OA) compared to that of the O mice, and for the young APP/PS1 (YA) mice compared to that of the OA group (**Fig.1 c**). The AUC for the spine was much lower than that of the NOB, but in the young mice it was 4.0-fold greater than that of the older mice (**Fig.1c-d**). The maximum intensity (at 150 mins) was similar for the Y, YA, and OA for the NOB region, but it was significantly lower in the O mice compared to the other groups (**Fig.1 e**). The maximum intensity was much lower for the spine compared to that of the NOB (**Fig.1 e-f**). The slopes for the steeper part of the curve were significantly lower in the O mice compared to the other groups (**Fig.1 g**). These data suggested that CSF clearance of tau was a slower process that that of BSA, as reported^24^.

**Fig.1.**
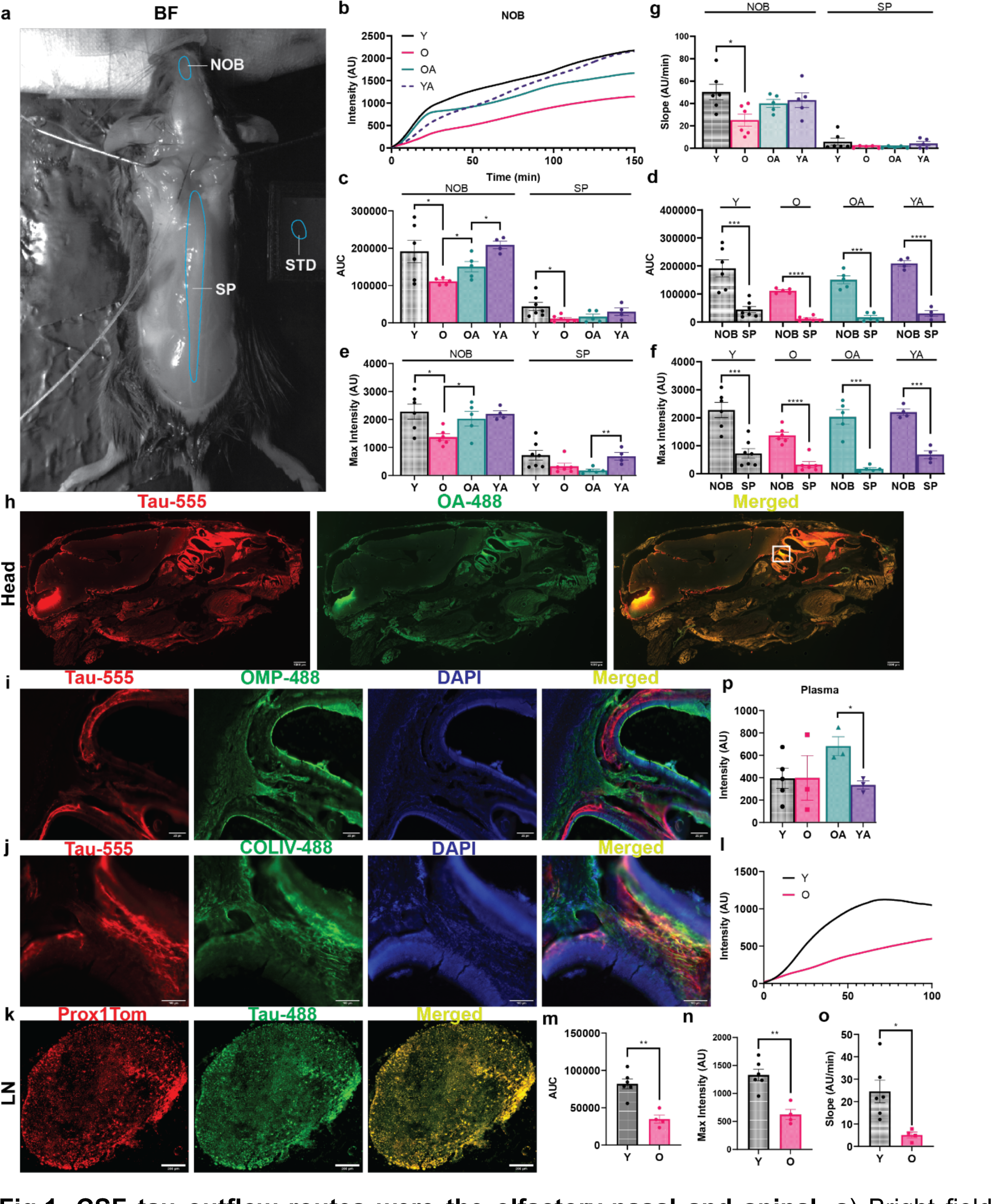
CSF tau outflow routes were the olfactory-nasal and spinal. **a)** Bright field image showing the mouse in its imaging position. Regions of interest (ROIs): Main cerebrospinal fluid (CSF) outflow regions: nasal/olfactory bulb (NOB) and spinal column (SP). STD, NIRF standard. **b**) Tau-NIRF intensity profile curves for the NOB regions for young mice (Y; black), older mice (O; red), and aged-matched young (YA; purple), and older (OA; blue) APP/PS1 mice. **c-g)** Area under the curve (AUC), maximum intensity at 150 mins and slopes (for the steeper part of the curve) for the NOB and spine in these mice. **h**) Representative sagittal section of the head after 150 mins showing tau-555 and OA-488 presence in the subarachnoid space and NOB regions in the older mice. **i**) Olfactory marker protein (OMP-647 but pseudo-color 488 nm) immunostaining for the region in the white box (cribriform plate region) in panel h. **j**) Collagen IV (COLIV-647 but pseudo-color 488 nm) immunostaining for the region of the white box in panel h. **k**) cervical lymph node of Prox1Tom reporter mouse showing the presence of tau-488 signal. **l-o**) Tau-NIRF intensity profile curves, AUC, maximum intensity, and slopes (for the linear part of the curve) for the superficial lymph nodes. **p**) Plasma tau-NIRF levels at 150 mins after CM injections. Values are mean ± SEM. Each solid circle was a mouse.

However, we confirmed that the nasal route across the cribriform plate was an outflow site for CSF tau-555 (**Fig.1 h**). While there was no significant colocalization of tau-555 with olfactory marker protein (OMP), a cytoplasmic olfactory receptor protein^54^, there was for collagen IV (a basement protein) along the tracks leaving the cribriform plate (**Fig.1 h-j**). In addition, we confirmed that the cervical lymph nodes were peripheral outflow sites for tau elimination from the CSF (**Fig.1 I**). Tau-NIRF clearance from the cervical lymph nodes (superficial) were reduced in the older mice compared to that of the younger ones (**Fig.4 l-o**). However, plasma tau-NIRF levels were unchanged between these two control groups, but it was significantly greater in the OA compared to that of the YA group at 150 mins after CM injections (**Fig.1p**).

Organs were removed at the end of the experiment (CM injection) and tau-NIRF intensity recorded. While signals were present on both the ventral and dorsal surfaces of the brain, there was little for the isolated skull cap, which contains the intact dorsal dura and dorsal skull bone (**Fig.2a**). NIRF-tau intensity was significantly greater on the ventral brain surface compared to that of the dorsal for the O, OA, and YA group, but not for the Y mice at 30 mins (**Fig.2b**). The average of the dorsal and ventral tau-NIRF intensities were then used. There were no differences between tau-NIRF brain and spinal uptake, except in the OA mice (**Fig.2c**) at 30 mins. There was a significant greater tau-NIRF uptake by the brain (BR) for the OA mice compared to that of the controls and YA group at 30 mins (**Fig.2d**). For the spine, the uptake was greater for the OA group compared to that of the YA group (**Fig.2d**). However, the differences between the tau-NIRF intensities between the dorsal and ventral brain surfaces were greater for the Y, OA, and YA but not for the Y group at 150 mins (**Fig.2 e**). In addition, these differences were greater at 150 mins compared to that at 30 mins (**Fig. 2b, e**). The brain tau-NIRF intensities were significantly greater for the O and OA compared to that of the Y and O or YA groups, respectively, using the average of the dorsal and ventral tau-NIRF intensities (**Fig.2f**). Brain tau-NIRF intensity was 1.7-to 2.3-fold greater at 30 mins compared to that at 150 min (**Fig.2 g**). CM injected tau-NIRF was also associated with the pial vessels (**Fig.2h**), and colocalized with PDGFRβ-positive cells (pericytes, **Fig.2i**). Thus, there was perivascular flow of CSF containing tau along the pial vessels, as reported by us and many others^51,55^.

**Fig.2.**
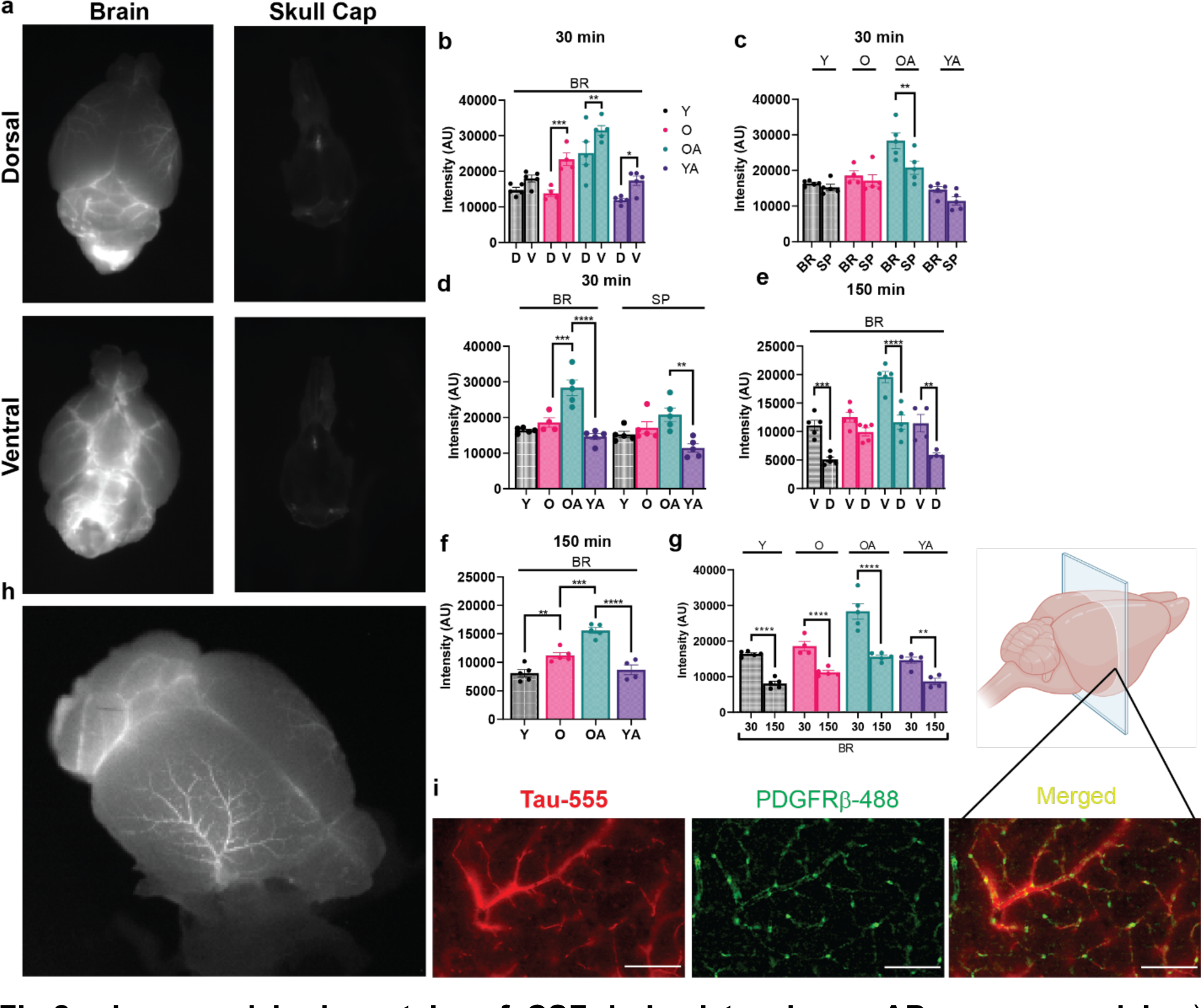
Increased brain uptake of CSF-derived tau in an AD mouse model. **a)** Representative tau-NIRF images on the dorsal and ventral brain and skull cap surfaces after 150 mins in young mice. **b**) Tau-NIRF brain uptake on the ventral and dorsal surfaces for young (Y), middle-aged (O) control mice, and aged-matched Y and O APP/PS1 after 30 mins. **c**) Average tau-NIRF intensity (D and V) for brain (BR) and spine (SP) at 30 mins. **d**) Comparison of the averaged tau-NIRF intensity for brain (BR) and spine (SP) at 30 mins for these groups of mice. **e**) Tau-NIRF brain uptake on the ventral and dorsal surfaces after 150 mins. **f**) Average tau-NIRF intensity (D and V) for brain at 150 mins in these groups of mice. **g**) Tau-NIRF brain uptake at 30 and 150 mins. **h**) Tau-NIRF uptake by the pial vessels on the brain surface at 150 mins. **i**) Tau-555 colocalized with platelet-derived growth factor receptor ý (PDGFRý)-positive cells (pericytes) in brain sections at 150 mins. PDGFRβ-647 was used but pseudo-color for 488 nm. Values are mean ± SEM. Each solid circle was a mouse. Scale bar = 100 μm.

CM injected tau-555 was detected in the parenchyma but mainly in the older and APP/PS1 mice (**Fig.3a-d**), and associated with NeuN-positive cells (neurons, **Fig.3c-d**) due to its neuronal uptake, as reported^56^. The injected tau-555 was colocalized with anti-tau antibody (T12), suggesting that there was intact human tau protein (**Fig.3d**). Tau-555 was present in the hippocampus and corpus callosum regions (**Fig.3d**). For the spinal cord, the injected tau-555 was present within the subarachnoid space (SAS) and spinal cord (**Fig.3e**).

**Fig.3.**
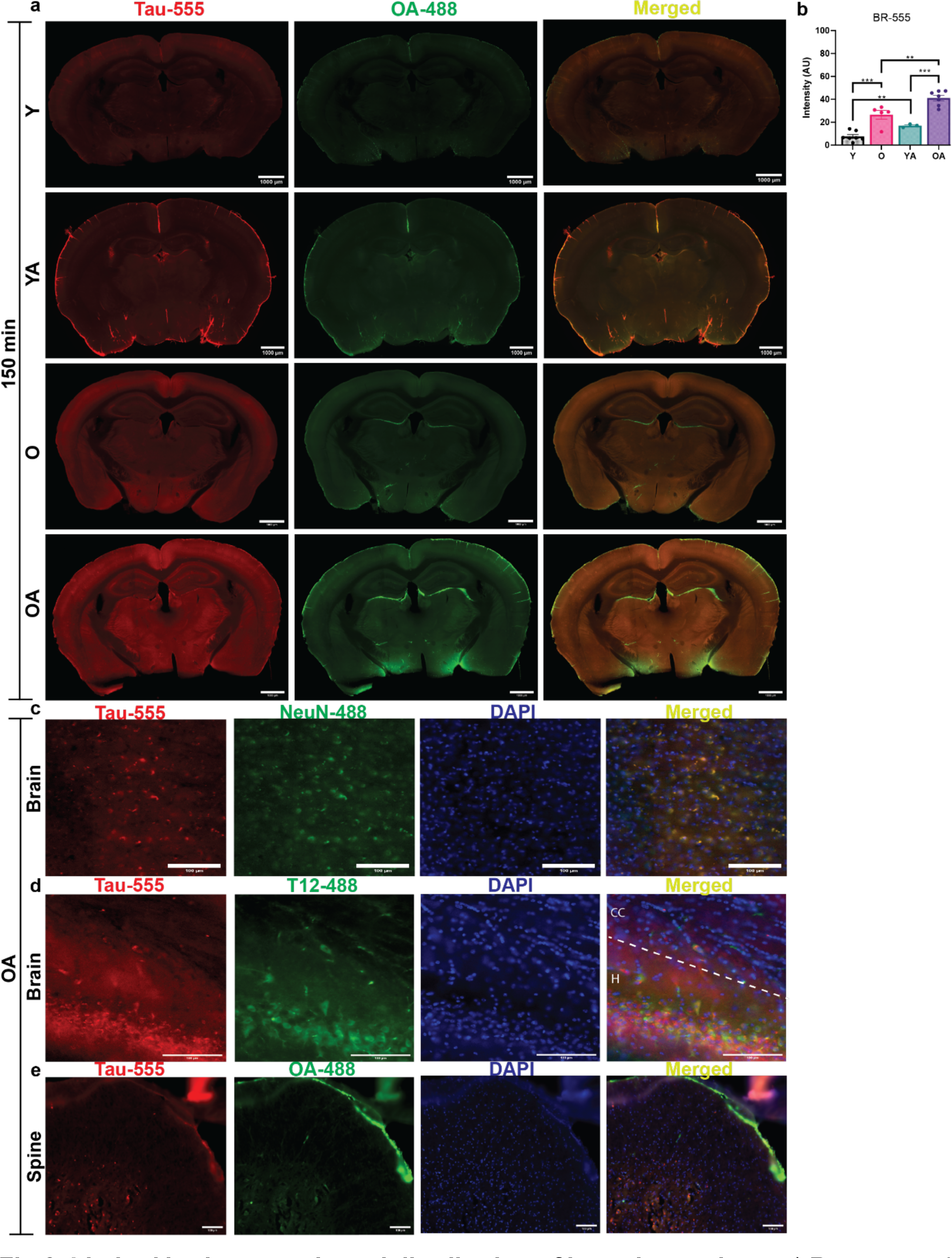
Limited brain parenchymal distribution of intracisternal tau. **a)** Representative coronal brain sections of young (Y) and middle-aged (O) control mice and young (YA) and middle-aged APP/PS1 (OA; blue) mice after 150 mins. **b**) Quantification of the average tau-555 intensity from the brain sections. **c**), Tau-555 colocalized with NeuN-positive cells (neurons). NeuN-647 was used but pseudo-color for 488 nm. **d**), T12 (anti-tau antibody) colocalized with injected tau-555 and was present in the hippocampus (H) and corpus callosum (CC) regions. T12-647 was used but pseudo-color for 488 nm. **e**), Injected tau-555 was present in the spinal cord of the older mice. Tau-555 and OA-488 were co-injected into the cisterna magna. Each solid circle was a mouse. DAPI (neuronal nucleus). Values are mean ± SEM. Scale bar = 100 μm.

CM injected tau-NIRF was distributed to many peripheral organs, including the liver, kidney, and bone (**Fig.4a**). The signal was more intense in liver even after 150 mins (**Fig.4a**). The femur NIRF-tau levels were increased over time (**Fig.4b**). In the whole organ tau-NIRF intensities were 5- to 10-fold greater for the liver and kidney compared to that of bone (femur) in young mice at 30 mins (**Fig.4c**). However, tau-NIRF uptake was significantly greater in the OA compared to that of the YA group for the liver, and in the O and OA compared to that of the Y and YA, respectively, in the kidney (**Fig.4d**). Tau-NIRF uptake in the OA was greater compared to that of the Y and O, respectively, for the femur (**Fig.4d**). **Fig.4e** shows a schematic diagram of CSF-derived tau protein distribution and retention within brain and to peripheral organs, including bone.

**Fig.4.**
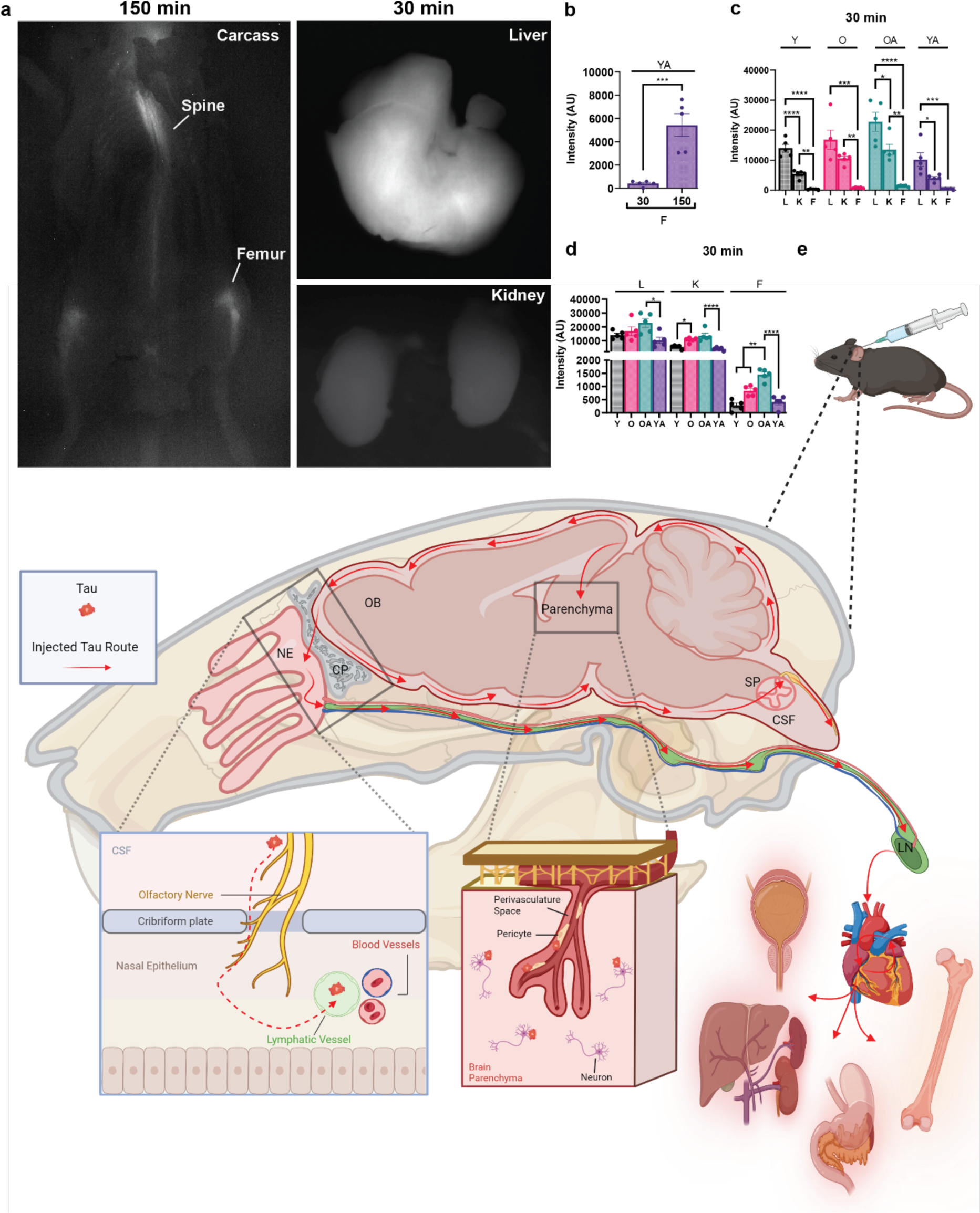
CSF-derived tau distribution to peripheral organs includes the skeleton. **a)** Representative tau-NIRF images for the skeleton, liver, and kidney. **d**) Tau-NIRF intensity at 30 and 150 mins in the femur after tau-NIRF intracisternal injection. **c-d**) Tau-NIRF intensity for the liver (L), kidney (K) and femur (F) at 30 mins. **e**) Schematic diagram of CSF-derived tau protein distribution and retention within brain, and to peripheral organs, including skeleton, liver, and kidney. CSF-derived tau drains via across the cribriform plate, and possibly spinal routes, into blood and retained in organs, such as bone. It was also excreted via the bladder. Values are mean ± SEM. Each solid circle was a mouse.

### Blood-derived tau multiorgan distribution includes the skeleton

To better assess tau’s peripheral distribution to bone, we recorded in vivo real-time uptake of blood-derived tau-NIRF in bone and compared it to that of liver, using a minimal invasive approach, whereby tau-NIRF was intravenously injected into the mice Thus, tau-NIRF signal was recorded with the mouse in its supine position and in organs/regions that were visible to the camera. The regions showing the presence of tau-NIRF signal were identified as ROIs (**Fig.5 a**). Tau-NIRF signal was predominantly in the liver, but there was distribution to bone, such as femur/knee and sternum (**Fig.5b-c**). There was a progressive increase in tau-NIRF uptake by these organs over the 150 mins, which approached a plateau from about 100 mins (**Fig.5b-c**). The area under the AUC for the liver was significantly greater by 6.0-8.0-fold compared to that of the femur/knee (F) and sternum (St) in the three groups of mice (**Fig.5 d-e**). In addition, the AUC of the liver and femur was significantly greater in the APP/PS1 (OA) mice compared to that of age-matched controls (**Fig.5d).** Similarly, the plateau intensity for the liver was significantly greater than that of femur and sternum in the three groups of mice (**Fig.5f-g**). In the APP/PS1 mice, the plateau intensity of the liver and femur was significantly greater than that of the age-matched controls by 1.8 to 2.5-fold (**Fig.5f**). The plateau intensity of the femur in the young mice was significantly greater by 2.0-fold compared to that of the older counterpart (**Fig.5f**). The rate of rise (slope) of the tau-NIRF intensity for the steep part of the curve was significantly greater, by 6.0 to 8.0-fold, for the liver compared to that of the sternum and femur in the three groups of mice (**Fig.5 h-i**). For the APP/PS1 mice, the slopes for the liver and femur were significantly greater compared to that of age-matched controls (**Fig. 5 h**). Thus, like the liver, tau-NIRF was retained in bone, at least up to the duration of the experiments (150 mins). In APP/PS1 mice, the uptake was greater for the liver and femur compared to controls.

**Fig.5.**
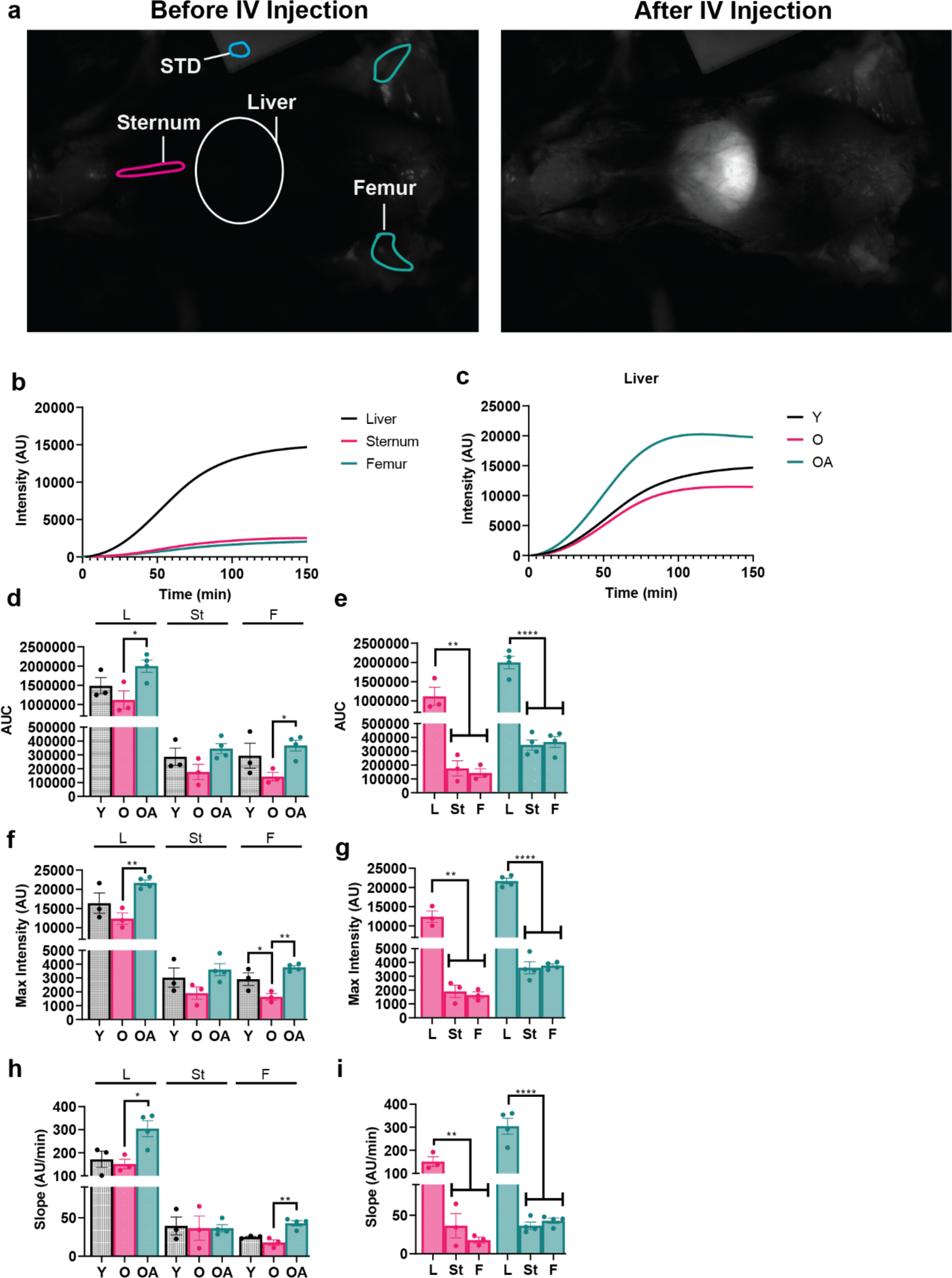
Kinetics of in vivo skeletal and liver distribution of tau protein. **a)** Representative images showing tau-NIRF distribution to liver, femur/knee, and sternum. **b**), Tau-NIRF distribution profile for the liver, sternum, and femur in young mature mice. **c**), Tau-NIRF distribution profile for the liver in young mice (Y; black), middle aged mice (O; red) and middle-aged APP/PS1 (OA; blue). **d-e**) Area under the curve (AUC) for the liver, sternum, and femur in these groups of mice. **f-g**) Maximum intensity of tau-NIRF for the liver, sternum, and femur for these groups of mice. **h-i**) Slope of the steep part of the tau-NIRF intensity profile curve for the liver, sternum, and femur for these groups of mice. Values are mean ± SEM. Each solid circle was a mouse. STD (NIRF standard block).

At the end of the experiment the mice were PFA fixed. The organs with detectable tau-NIRF signals were removed and imaged using the same parameters. An unexpected striking feature was the presence tau-NIRF signals in bone, including the spine, cranium (skull cap) on both the ventral and dorsal surfaces, and in the femur (**Fig.6a-d**). Also, signal was detected on the ventral surface of the brain especially at the region of the pituitary gland/median eminence (PG) and on the dorsal surface of the head (**Fig.6 e-f**). Tau-NIRF signal was prominent in the liver, kidney, and in the intestine (mainly in the duodenum), which was mostly within the lumen since removing its content dramatically reduced tau-NIRF signal (**Fig.6 g-j**). Signal was present in the gall bladder (**Fig.6 g**). Tau-NIRF signal was also present in the spleen and urinary bladder (**Fig.6 k-l**). Tau-NIRF signal was also present in the heart (**Fig. S1**). Thus, in addition to the expected tau distribution to the liver and kidney, tau was present in bone, the intestine and gall bladder. These images suggested that the monomeric form of full-length tau (∼45kDa) can be eliminated via the urinary bladder and possibly the intestine. Also, the data suggested that tau was accumulated in many organs, including the skeletal bone. It also suggested that there were significant levels of blood-derived tau in the isolated skull cap, which was like that of the head (intact brain and skull cap). Brain tau levels was low, and better resolved by imaging it isolated from the skull.

**Fig.6.**
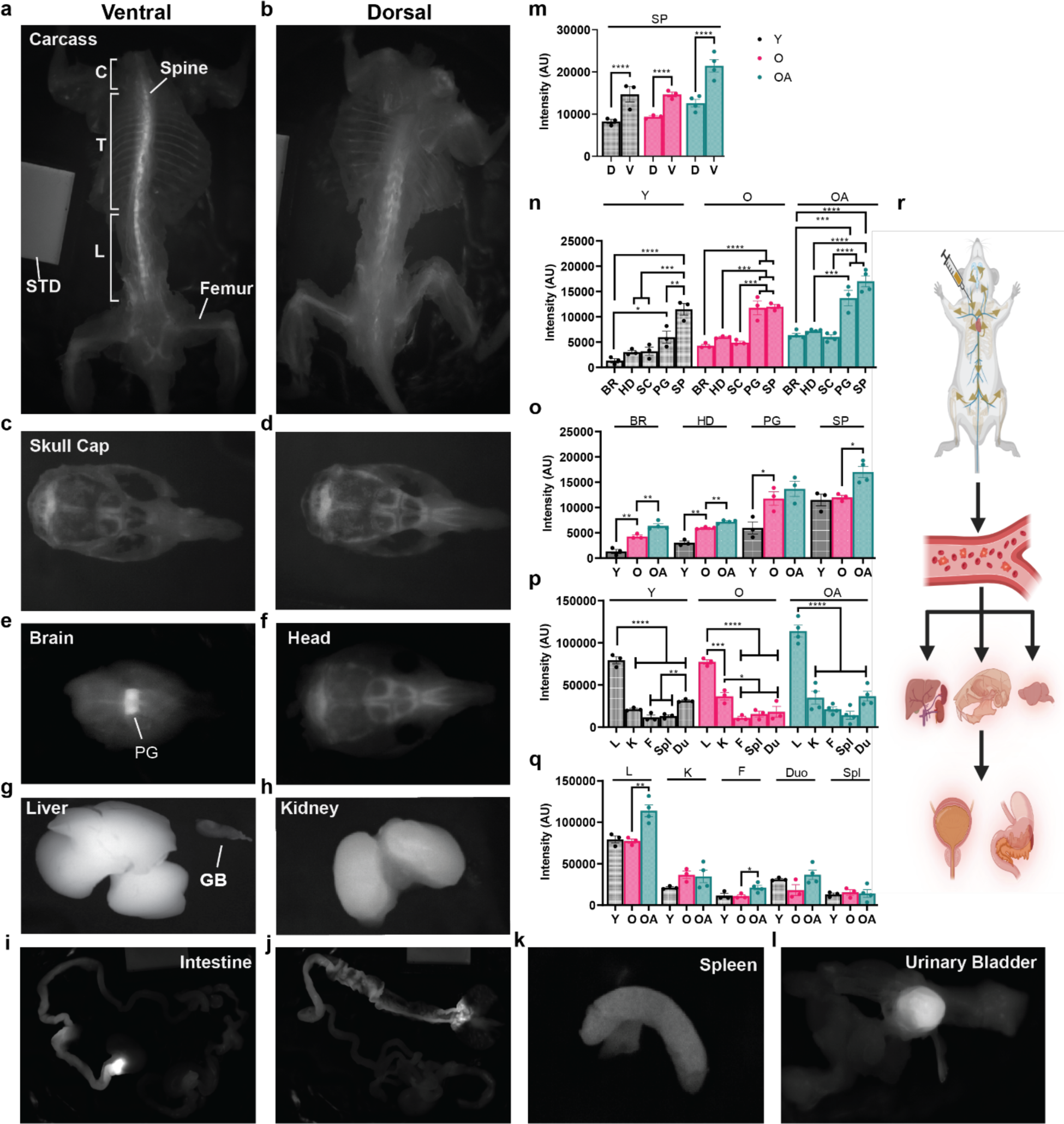
Multiorgan distribution of blood-derived tau includes the skeleton. **a-l)** Representative images for the ventral and dorsal skeleton (**a-b**), skull cap (**c-d**), brain (**e**), head (**f**), liver and gall bladder (**g**), kidney (**h**), intestine (**i-j**), spleen (**k**), and urinary bladder (**l**) at 150 mins after IV tau-NIRF. **m**) Tau-NIRF spinal uptake by the dorsal and ventral surfaces for young (Y; grey) and middle-aged (O; red) control mice and middle-aged APP/PS1 (OA; blue) mice. **n-o**) Tau-NIRF uptake by the brain (BR), head (HD) skull cap (SC), base of brain at the pituitary gland area (PG), and spine (SP) using mean levels for the ventral and dorsal surfaces. **p-q**) Tau-NIRF uptake by the liver (L), kidney (K), femur (F), spleen (Sp), and duodenum (Du). **r**) Schematic diagram showing multiorgan organ distribution of blood-derived tau and possible excretion routes. Values are mean ± SEM. Each solid circle was a male mouse. STD (NIRF standard block).

The tau-NIRF intensity was greater on the ventral surface of the spine, by 1.5-to 1.8-fold, compared to that of the dorsal surface in young mature (Y), middle-aged controls (O) and middle-aged APP/PS1(OA) mice (**Fig.6 m**). Using the average of the dorsal and ventral tau-NIRF intensities, the spinal signals were 2.0 to 4.0-fold greater than that of the head in these groups of mice (**Fig.6n**). Tau-NIRF intensities were the lowest for the head/brain compared to that of PG region and the spine (**Fig.6n**). Tau-NIRF signals for the brain (BR), head (HD) and PG were 2.5- to 4.0-fold greater in the O group compared to that of the Y group of mice, while there were no significant differences for the spine (**Fig.6 o**). However, tau-NIRF signals were significantly greater in the OA mice compared to that of the O for the BR, HD, and SP but not for the PG (**Fig.6 o**). Thus, some blood-derived tau was associated with brain, and this was greater with aging and in the APP/PS1 mice. For the peripheral organs, tau-NIRF intensity for the liver (L) were 5.0- to 8.0-fold greater than that of duodenum (Du), kidney (K), spleen (Sp) and femur (F) (**Fig.6 p**). However, there was no significant differences for these organs/regions in these groups of mice, except for the higher levels in the liver and femur in the OA compared to that of the O group (**Fig.6 q**). Thus, there was differential uptake of blood-derived tau for the peripheral organs, but it was greater in the liver and femur in the OA mice compared to that of the O group (**Fig.6r**).

We explored possible relevance of tau protein in bone by immunostaining for integrin-binding sialoprotein (IBSP), a major non-collagenous glycoprotein that is abundantly expressed by osteoblast and associated with mineralization^53^. Injected tau-555 was colocalized with IBSP in the femur (**Fig.7a-d**). In the trabecular bone tau-555 colocalized with IBSP (**Fig.7 b**). This was confirmed by correlation analysis (**Fig.7c**). In addition, tau-555 was colocalized with IBSP in the bone marrow (**Fig.7 d**). We then used the Tg-htau-GFP transgenic mice, which express htau but not the mouse’s tau, and show that there was htau in brain (as expected), femur, and femur head **(Fig.S2a-b**). Similarly, h-tau was present in the liver (**Fig.S2c).**

**Fig.7.**
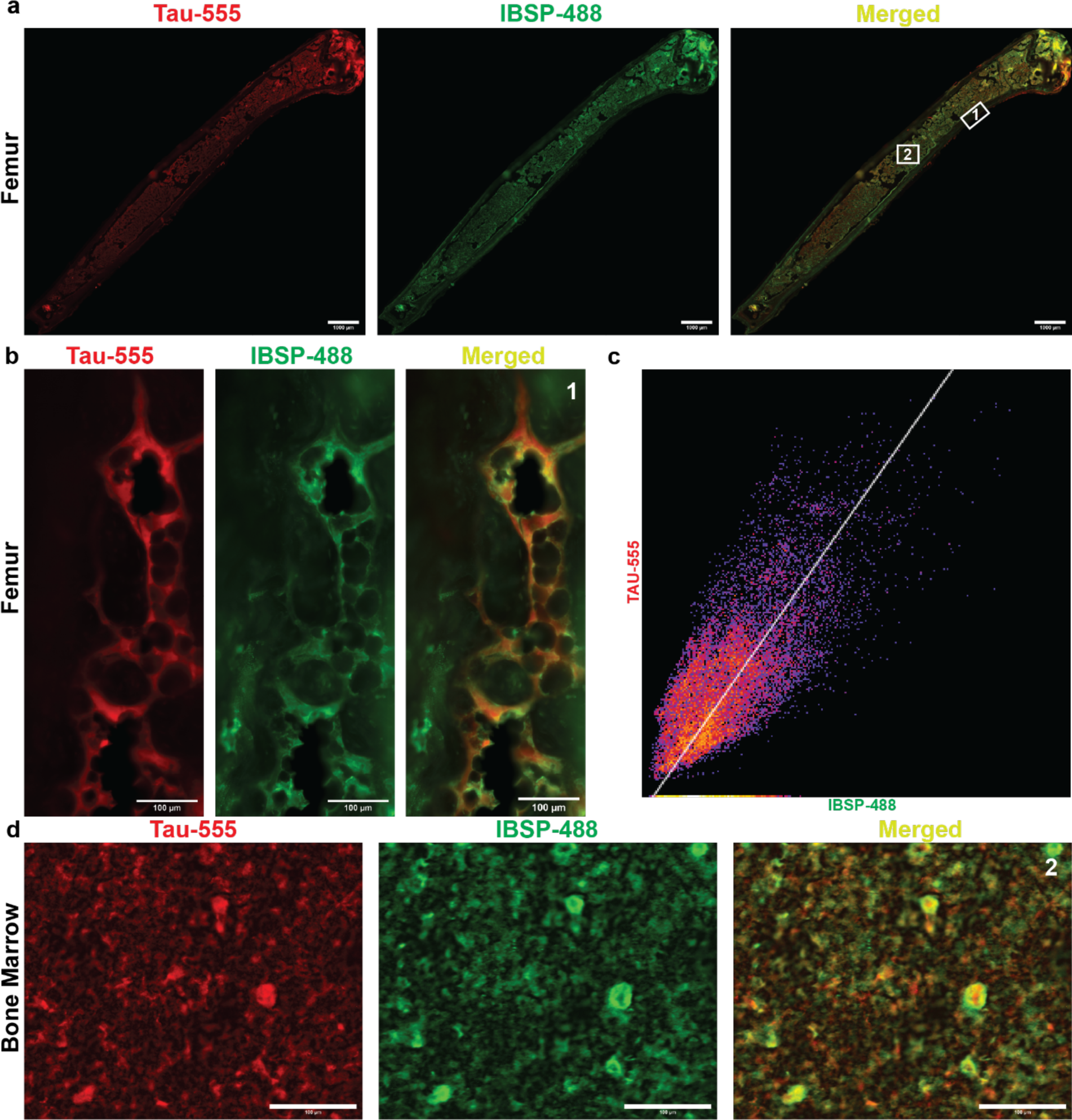
Blood-derived tau colocalized with integrin-binding sialoprotein (IBSP) in bone. **a**) Representative images showing tau-555, IBSP-488 and merged image for the femur. **b**), Tau-555 colocalized with IBSP in the trabecular bone (white box 1). **c**), Correlation between tau-555 and IBSP-488 in the trabecular bone. **d**), Tau-555 colocalized with IBSP in the bone marrow (white box 2). N= 3. Scale bar = 1000 μm for panel a and 100 μm for panels b-d. IBSP-647 was used but was pseudo-color IBSP-488 for the presentation.

### Anti-tau antibody increased CSF tau clearance

Since tau protein was retained in brain and in peripheral organs, we explored potential therapeutic intervention to promote clearance using a commercially available anti-tau antibody (T12). Anti-tau antibodies have been used to alleviate tau’s toxic effects^57,58^. Tau-NIRF/anti-tau complex (tau-NIRF-AB) was prepared and the same dose of tau-NIRF was used as in the control. In contrast to controls, there was a rapid rise in tau-NIRF-AB signal at the NOB site, which peaked at about 30 mins, followed by an exponential fall (**Fig.8a**). This pattern of the intensity profile curve was like that of albumin clearance from the CSF^24^. At the NOB, the AUC, maximum intensity, and rate of rise (slope) were increased significantly (by 2.0 to 6.0-fold) for the complex compared to that of controls tau-NIRF(**Fig.8b-d**). The data suggested that T12 complexed to tau likely prevented tau retention, and thus, promoted tau elimination. This may explain the even lower levels of the complexed tau in brain compared to that of controls (**Fig.8e-g**).

**Fig.8.**
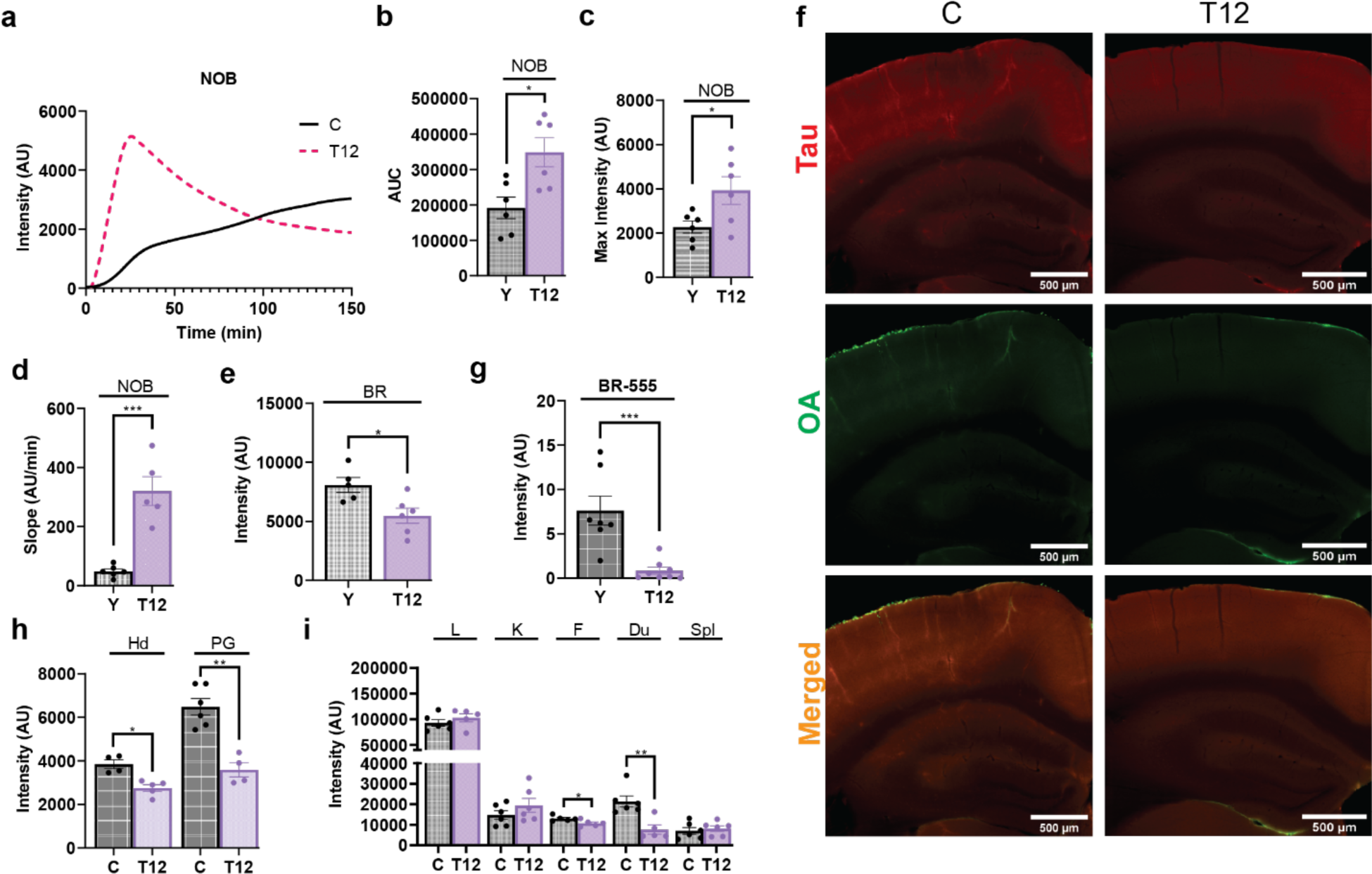
Anti-tau antibody increased CSF tau outflow but differentially affected its peripheral distribution. **a)** Tau-NIRF intensity profile curves for the NOB regions for young mice with and without anti-tau antibody (T12) after its intracisternal injection. **b-d)** Area under the curve (AUC), maximum intensity and slopes (initial linear part of the curve) for the NOB in these mice. **e**) Brain tau-NIRF uptake. **f**) Representative coronal brain sections after 150 mins showing little tau-555 and OA-488. **g**) Quantification of the average tau-555 intensity from the brain sections. **h**) Tau-NIRF intensity of the head (Hd) and base of brain at the pituitary gland area (PG). **i**) Tau-NIRF intensity of the liver (L), kidney (K), femur (F), duodenum (Du) and spleen (Spl). Values are mean ± SEM. Each solid circle is a mouse. Scale bar = 100 μm.

### Blood anti-tau antibody differentially affected the distribution of blood tau

Since NIRF-tau uptake in the liver and bone appears to approach equilibrium after about 100 mins, experiments were performed at 60 min after IV injections. The tau-NIRF-AB complex or tau-NIRF was injected IV via the jugular vein, and after 60 mins the mice were PFA perfused, and organs removed and imaged. The complex significantly decreased tau uptake into the pituitary gland region and head (**Fig.8h**). For the peripheral regions, the complex reduced tau-NIRF uptake into the femur and duodenum but not into liver, kidney, and spleen (**Fig.8i**). While further work is needed on the dose and duration, these pre-clinical data show promise.

## Discussion

In these studies, we explore tau distribution to peripheral organs and within brain in controls and APP/PS1 mice at two age groups and tested the feasibility of an intervention to promote tau clearance. Our main findings were as follows: **1**. While CSF- or blood-derived tau protein was distributed and accumulated in many peripheral organs, including the liver, skeleton, kidney, intestine, spleen and both gall and urinary bladders, it was more pronounced in the liver. **2**. Spinal tau accumulation was intense, and more prominent on the ventral surface. **3**. In the femur, the administered tau was colocalized with integrin-binding sialoprotein (IBSP), a major non-collagenous glycoprotein that is associated with mineralization. **4**. Age differentially affected the distribution of blood-derived tau. **5**. In the older APP/PS1 mice, tau accumulation was increased in liver, brain, spine, and femur compared to controls. **6**. The rate of tau accumulation in liver was greater than that of femur, and greater in APP/PS1 mice for both liver and femur compared with that of controls. **7**. Tau was mainly cleared slowly from CSF via across the cribriform plate (NOB) and cervical lymph nodes since there was no falling phase from the intensity prolife at the NOB up to 150 mins. CSF-derived tau was present in blood. The rate of tau appearance at the NOB was greater in young mice compared to the older counterpart, and APP/PS1 mice. **8**. In general, inflow of CSF tau into brain was low, but greater in the older and APP/PS1 mice than controls. CSF tau was associated with NeuN-positive cells and PDGFRβ-positive cells. **9**. Anti-tau antibody in the CSF considerably increased the CSF tau clearance, while blood anti-tau antibody decreased tau accumulation in the femur but not in liver, kidney, and spleen. **10**. The data show that CSF-derived tau was present in blood, which was distributed and accumulated in many organs, including bone, and excreted by the kidneys, and possibly the intestine (via bile). Anti-tau antibody increased the clearance of CSF-derived tau protein, and differentially affected its accumulation in peripheral organs. The significance of tau accumulation in bone, and each of the other organs need further work.

The striking accumulation of CSF-derived tau by the skeleton was surprising and raises the tantalizing questions about its importance. Bone receives about 5-15% of the cardiac output, and blood flow is heterogenous and affected by its function^59–63^. Bone capillary should be permeably to the monomeric form of tau protein (∼45 kDa), since albumin (∼66 kDa) enter bone ISF from blood^64,65^. However, the tau-NIRF intensity profile for bone (femur) suggested that it’s accumulated over the duration of the experiment (150 mins). The accumulation of tau in bone appears to be selective, as tau protein was colocalized with integrin bone sialoprotein (IBSP). IBSP, a major member of the SIBLING (small integrin-binding ligands N-linked glycoprotein) family of non-collagenous extracellular proteins, is produced by osteoblasts, osteoclasts and osteocytes, binds to integrins and involved with mineralization^53,66–71^. Also, IBSP is involved in the mediation of endothelial cell attachment, migration, and angiogenesis^72–74^. The endothelium, especially the subgroup that highly expresses CD31 and Endomucin, appears to be central to the formation of bone by forming a local niche microenvironment^75,76^. It is unclear on whether CSF-derived tau proteins accumulation in bone is associated with bone function or degeneration. However, cardiovascular diseases (SVD) are associated with osteoporosis and loss of bone density is associated with calcification of the vasculature^77–83^. CVD and tau proteins are also associated with neurodegeneration^84–88^. In the middle-aged APP/PS1 mice, there was greater accumulation of blood/CSF-derived tau protein in the skeleton (spine and femur) compared to aged-matched controls. Tau uptake was also increased in the skeleton with aging. Is this a possible brain to skeleton link, and a contributing factor in degeneration? We did not use tau models of AD to avoid possible interaction between endogenous and injected tau.

CSF-derived tau-NIRF distribution to the skull was low but was more prominent with blood-derived tau-NIRF. The peripheral distribution of CSF-derived tau-NIRF to the skull from blood will be limited by the greater distribution to tissues, such as liver and other bones, and likely due to the differences in blood flow. A significant flow of CSF-derived albumin directly to the cranium was not detected^24^. Similar data were obtained for other CSF-derived molecules^89^.Others have suggested that CSF flow to the dura/cranium^90–93^. Our data suggested that blood-derived tau is distributed to the cranium, like other skeletal structures, and this likely limited by blood flow. CSF-derived tau in blood is also distributed to the cranium, but this is reduced by the slow outflow of CSF into blood and the rapid distribution by blood to all organs.

While the accumulation of brain-derived tau in liver may not be surprising, its presence in the gall bladder and the lumen of the intestine was, which may indicate a pathway for its elimination, and/or absorption via the enterohepatic circulation. In contrast to SARS-CoV-2/SP, a protein of similar molecular weight, which reached a peak in liver a few minutes after IV injection and then rapidly fell, tau protein was accumulated over the 150 mins^44^. This would suggest a specific tau protein uptake mechanism for its accumulation in liver, and possibly its secretion with bile^94^. Bile is released into the duodenum of the small intestine, which is where blood-derived tau protein was present. Bile secretion is variable and any excretion of tau protein via this pathway will also be variable^94^. The presence of tau protein in the kidney and urinary bladder suggested a possible elimination route for tau protein. However, tau protein accumulation in the urinary bladder will be variable since urination was not controlled for. While tau protein (∼45 kDa) will be filtered by the kidney, it is unclear on whether there is reabsorption. Further work is needed on how the liver and kidney process CSF-derived tau proteins to understand their role in controlling the levels of tau proteins in blood.

Levels of tau proteins in CSF are used for diagnosis of dementias, and bulk flow of CSF is a pathway for the clearance of tau proteins into blood^95–97^. The CSF outflow sites of tau protein after intracisternal injection was as reported for albumin^24^. These outflow sites were mainly across the cribriform plate (nasal/olfactory route) and possibly spinal routes, as reported by us and others^24,89,98–105^. However, at the NOB, the tau-NIRF intensity profile did not peak over the 150 minutes, while that of albumin peaked at around 30 minutes^24^. There was a rise in tau-NIRF intensity to a deflection point at about 30 minutes followed by a progressive slower increase. This suggested that tau protein is retained, and there is a net uptake at the ROI, while there is some appearance of tau at the cervical lymph nodes. Aging reduced the initial rate of rise (appearance) of tau-NIRF intensity at the NOB, and its elimination via the cervical lymph nodes^24^. The initial slope for albumin was about 10 times greater than that of tau^24^. This suggested that less of the tau protein arrived and/or remained at the ROI possibly due, perhaps, to the reduced CSF secretion with aging, a driving force for the flow of CSF^106–108^, and retention by binding to tissues. Concomitant with this, there is greater retention within the brain parenchyma in the older mice compared to the younger ones, due to greater diffusion and/or perivascular inflow^51,109^. Unexpectedly, the tau-NIRF intensity profile graph for the APP/PS1 mice was between that of the young and older control mice. This could be due to the prominent presence of amyloid-β pathology and gliosis from about 2 months of age^110^. While the CSF levels of murine tau protein is increased with aging in the APP/PS1 mice compared to controls, the relevance is unclear^111^.

CSF-derived tau-NIRF and tau-555 were associated with pial vessels, and colocalized with PDGFRβ-positive cells, especially in the older mice. Similarly, there was colocalization of CSF-derived tau-555 with NeuN-positive cells, which was likely due to tau uptake in neurons^56^. There was some tau-555 within the corpus collosum, a likely low resistance pathway for the parenchyma flow of ISF^112,113^. Parenchymal distribution of CSF-derived tau was limited by the flow of CSF via the cribriform plate, as reported^24,89^. In general, inflow of CSF-derived tau into brain was low but increased in the older mice and in the APP/PS1 mice.

A major CSF peripheral outflow site is via the cribriform plate, and the cervical lymph nodes and other lymph nodes^24,101^. Aging reduced the clearance of tau-NIRF via the cervical lymph nodes^24^. There are reports of reduced clearance of CSF-derived molecules via the peripheral lymphatic system^24,108^. The flow of CSF-derived tau along the entire spine was low compared to that of the cranium, which explains the absence of detectable tau-NIRF in the sacral lymph nodes^101^. Plasma levels of CSF-derived tau-NIRF, at 150 minutes, was greater in the older APP/PS1 mice compared with controls.

It was shown that there is a high residual serum level of radio-iodinated full-length tau (^125^I-tau) after its IV injection^114^. This was about 25-30% of the injected dose (∼1.5 (log %injected dose/ml)), which suggests that there was significant sequestration (binding) of tau by serum molecules, such as proteins. Binding of tau to plasma molecules would limit its plasma clearance^115,116^. This would explain why there is a progressive increase in tau-NIRF intensity in organs. It is unclear on why plasma levels tau was greater in the older APP/PS1 mice. Further work is needed on tau binding molecules in plasma, and if they are changes with aging and in AD.

Anti-tau antibody has been shown to reduce soluble tau oligomer effects^57,117^. Anti-tau antibody (T12) complexed to tau-NIRF (T12-tau-NIRF) increased tau-NIRF clearance at the NOB region, to a pattern like that obtained for albumin^24^. This faster tau-NIRF clearance at the NOB reduced its inflow into brain, which supports the concept that faster clearance at the NOB region reduced CSF distribution to brain parenchyma. Intravenous injection of the complex reduced tau-NIRF distribution to the head, PG region, femur, and duodenum, but not for the other organs, such as liver, kidney, and spleen. These differential effects of T12 on the distribution of tau-NIRF suggest that the free and complexed tau is distributed to organs differently. Further work on mechanism of tau entry into each organ, such as whether there is receptor mediated or adsorptive endocytosis, is needed to be able to provide a meaningful explanation.

## Summary

Tau protein clearance from the CSF was slower than albumin, but T12 increased it to a pattern like that of albumin. While the main CSF outflow pathways were like that of albumin, there was some tau protein accumulation within the brain parenchyma that was associated with neurons and pericytes. Blood-/CSF-derived tau was retained in organs, such as brain, skeleton (femur), and liver, which was aggravated with aging and in a mouse model of AD (APP/PS1). In the femur, injected tau colocalized with IBSP, but the significance needed further work. Since tau protein was present in the lumen of the intestine and in the urinary bladder these were possible excretory routes. There is blood-derived tau protein associated with brain, a possible re-entry route, which was greater with aging and in the APP/PS1 mice. CSF anti-tau antibody (T12) increased tau protein clearance at the nasal/olfactory bulb region, and reduced tau protein inflow/diffusion into brain. Blood T12 reduced brain tau levels, and uptake into the femur and duodenum, but not in liver, kidney, and spleen. The progressive accumulation of tau in various organs suggests that blood tau levels are not rapidly cleared. It is likely that tau is sequestered by plasma molecules. While the data suggest that tau may have effects beyond brain, further work is needed on the mechanism of tau uptake in organs, such as bone, liver, kidney, to better understand the relevance to tauopathy. Similarly, while the data suggest that blood tau is not rapidly eliminated, further work is needed on the sequestration molecules in blood, to better understand the role of plasma/serum tau levels in tauopathy.

## Materials and Methods

### Materials

Full length tau monomeric protein form (2N4R) was obtained from Dr. Kayed Lab and stored at -80°C^45–47^. Tau was labeled with near infrared fluorescent molecule (NIRF) by using a kit (IRDye800CW Microscale kit, Li-COR Biosciences, Nebraska, USA), and by following the manufacturer instructions. This conjugation forms a stable amide bond (Li-COR Biosciences). Tau protein was also labeled separately with Alexa Fluor 555 using a kit (Microscale protein labeling kit; ThermoFisher Scientific; Waltham, MA, USA) and by following the manufacturer instructions. In addition, both labels were purified using 3 kDa molecular weight cut-off ultrafiltration filter (Amicon Ultra Centrifugal Filter, Millipore). There was no detectable dye in the filtrate for samples. Ovalbumin (OA) Alexa Fluor-488 conjugate (molecular weight 45 kDa) were obtained from Life Technologies Corporation, Carlsbad CA, USA.

### Mice

C57BL/6J (2–12 months old), APP/PS1 (B6.Cg-Tg(APPswe,PSEN1dE9)85Dbo/Mmjax (2-12months old; MMRC) and Prox1Tom reporter [B6;129S-Tg (Prox1-tdTomato)12Nrud/J (Prox1Tom)] at 12 months old were obtained from Jackson Laboratory (Bar Harbor, ME, USA). Tg-h mice (B6. Cg-*Mapt^tm^*^1^*^(EGFP)Klt^* Tg (MAPT)8cPdav/J) were obtained from Dr. Kayed Lab. Male mice were used. All animal studies were performed in accordance with the National Institute of Health guidelines and using protocols approved by the University Committee on Animal Resources. Mice were housed in the vivarium of the University of Rochester, School of Medicine, and Dentistry, on a 12:12 light/dark schedule (6:00 a.m.–6:00 p.m.) with food and water ad libitum.

### Intracisternal injections

Mice were anesthetized with isoflurane and placed in the prone position on the temperature-controlled stage (Indus Instruments, Webster, TX, USA) to maintain body temperature. The hair was removed with a depilatory cream before surgery. A skin incision was made in the midline from the nose to the end of the spinal cord and the skull and spine column exposed to enhance the NIRF signal intensity. The cervical spine was exposed by removing the fat tissues on the back of the neck. Local anesthetic was applied to the exposed areas. The cisterna magna was exposed, cannulated with a 30G needle, connected to a 10 µL Hamilton syringe and a pump (Infusion/Withdrawal Pump II Elite Programmable syringe pump, Harvard Apparatus, Massachusetts, USA), as reported^24,44^. The injection site was sealed with cyanoacrylate (All Purpose KrazyGlue, Elmer’s). Fluorescent tracers (5 μL tau-NIRF or tau-555; 0.2 μg μL^−1^) and a protein reference molecule of similar molecular weight (ovalbumin-488 (OA-488, 0.1%; dissolved in aCSF, pH 7.4) were intracisternally infused (0.5 μL/min).

### In vivo dynamic imaging

NIRF intensities were measured using a custom-made NIR system^24,44^. The imaging system was composed of a lens (Zoom 7000, Navitar, Rochester, NY, USA), a NIRF filter set (Semrock ICG-B, IDEX Health & Science LLC, Rochester, NY) and camera (Basler acA3088-57um USB 30, Basler, ID: 107402). NIRF was excited with a tungsten halogen bulb (IT 9596ER, Illumination Technologies, Inc., Syracuse, NY, USA) through a ring illuminator (Schott, Elmsford, NY, USA). Imaging settings and recordings were accomplished through a custom-built LabVIEW program (National Instruments, Austin, TX, USA). Real-time NIR imaging was performed before injections (background) and every 4 sec for 150 min after tau-NIRF injection into the cisterna magna (CM) or intravenously (IV)^24,44,50^. Using ImageJ software (National Institutes of Health, Bethesda, MD, USA), ROIs were identified and the fluorescence intensity recorded at the same settings, for CM and IV injections^24^. A standardized fluorescent block was used to ensure the camera setting was the same for the experiments. The person performing the imaging and analysis was blinded to the experimental design. Tau fluorescence signal was enhanced to better show details for the images in the figures, but not for quantification, where the same imaging settings were used.

### Intravenous injections

Mice were anesthetized with isoflurane since it has fewer systemic hemodynamic effects^24,44,48,49^. Anesthesia was maintained with 1–2% isoflurane in oxygen. The hair was removed with a depilatory cream before surgery. The anesthetized mouse was placed on a temperature-controlled stage to maintain body temperature, a midline incision was made from the neck to the pelvic and the skin retraced to enhance the NIRF signal intensity. Local anesthetic (Topical Lidocaine 4% gel, ESBA Laboratory, Inc., FL, USA) was applied to the exposed areas. The right external jugular vein was exposed for a minimal invasive intravenous injection (IV) using a 30 G needle connected to a 500 µL insulin syringe via a tubing. The tubing contained the fluorescent tracer (10 μL), which was tau conjugated to IRDye800CW (Tau-NIRF; 4.6 pmol μL^−1^ (0.20 μg μL^−1^)), or Tau conjugated to Alexa-555 (Tau-555; 0.20 μg μL^−1^) and a protein reference molecule of similar molecular weight (ovalbumin-488 (OA-488, 0.1%), dissolved in PBS (pH 7.4). The bolus injection contained 10 μL PBS, 1 μL air gap, 10 μL tracer, 1 μL air gap, and 50 μL PBS to wash in the tracers completely. The injection was completed in 1 min. We used NIRF to minimize the natural background fluorescence of biomolecules, increase the signal to noise, and to provide a better contrast between the target and background^24,44^. We also used tau-555 to visualize and map its distribution in tissue sections and compared this to that of a reference protein molecule of similar molecular weight (OA-488). The concentration of tau protein was like that used in an earlier report^114^. According to this report about 25% of the injected dose was the residual concentration (retained in plasma). Thus, an estimated steady-state concentration would be 500 pg/ml. The effect of tau concentration on its uptake is unclear since there is no detail study on the mechanisms of transport by each organ. If there is receptor/carrier mediated uptake of tau, then uptake will be limited by the maximum transport capacity (Vmax). On-the-other-hand, if there is simple diffusion, then uptake will be dependent on tau concentration difference.

### Tissue preparation for fluorescence imaging

After the duration of the experiment, mice were transcardially perfused using cold PBS and paraformaldehyde (PFA, 4% in PBS pH 7.3). Tissues samples were removed, stored overnight in PFA at 4 °C, and re-stored in cold PBS solution containing 30% sucrose. Brain was cut into 100 µm coronal sections using a vibratome (Leica VT1000E). Liver, and lymph nodes were embedded in Optimal Cutting Temperature compound and cut into 30 μm sections using a cryostat. In some experiments, the skin and muscle were removed from the head, spine, and femur, then decalcified in 13% EDTA for 5-7 days, and cut using a cryostat, as reported^44^. Sections were mounted on Superfrost Plus glass slides using ProLong Gold Antifade Mountant medium (ThermoFisher Scientific, Waltham, MA, USA) for fluorescence imaging (VS120 Virtual Slide Microscope, Olympus). For brain and femur sections, the magnification (10x), exposure (200 ms) and gain (0) were fixed for all experimental groups based on pilot experiments. For spine sections the magnification was set at 20x, and the other parameters were the same. All fluorescence (tau-555 and OA-488) quantification was performed without enhancement of signals. Tau-555 was used since we do not have access to a microscope to analyze tau-NIRF in tissue sections. The person performing the imaging and analysis were blinded to the experimental design and tracer used. This was finally decoded when the figures were prepared for publication.

### Immunohistochemistry

Brain sections were immunostained for neurons (NeuN), pericytes (PDGFRβ), and Tau 12 (T12), and femur sections for integrin binding sialoprotein (IBSP). Whole head sections were immunostained for olfactory marker protein (OMP) and collagen IV (COLIV). The tissue sections were washed three times and placed in a blocking buffer of 5% donkey serum for 1 hr at RT. Sections were than labeled with the primary antibodies and incubated overnight. The primary antibodies were chicken anti-NeuN (1: 100, Abcam. Cat# ab134014), rabbit anti-collagen IV (1: 100, Cat# SAB4500369, Millipore Sigma), rabbit anti-IBSP (1: 100; ThermoFisher Scientific. Cat# PA5-114915), goat anti-PDGF Rβ (1: 100, R&D Systems. Cat# AF1042), rabbit anti-OMP (1:100, Abcam. Cat# 183947), mouse anti-tau (T12, 1:100, BioLegend. Cat# 806501). Sections were washed three times, and the secondary antibodies used were donkey anti-chicken-647 (1:300, Jackson Immunoresearch labs. Cat# 703-175-155), donkey anti-rabbit-647 (1:300, ThermoFisher. Cat#A32795), donkey anti-goat-647 (1:300, ThermoFisher. Cat# A32849), donkey anti-mouse-647 (1:300, ThermoFisher. Cat# A32787), which were added and incubated for 1.5 h at RT. Immunofluorescence was visualized by using an epifluorescence microscope (VS120 Olympus). For presentation and analysis, the 647 nm was pseudo-color with 488 nm for better visualization.

### Ex vivo NRIF imaging of whole tissues

At the end of in vivo imaging, animals were PFA perfused, and the whole animal and its organs were preserved, including the head/brain, spinal column, liver, lungs, heart, kidneys, spleen, and intestine, and skeleton, as reported^44^. Tissues were washed equally and imaged individually on the ventral and dorsal surfaces using the same parameters as that used for the in vivo imaging. Average of the ventral and dorsal surfaces were the same, except for the head/brain and spine. The averaged tau-NIRF intensity was used for comparison.

### Ex vivo fluorescence imaging of tissue sections

Ex vivo epifluorescence microscopy was used to evaluate the degree of tracer distribution in the tissue sections. Exposure and gain were fixed for all experimental groups based on the pilot experiments. To quantify the extent of tracer intensity distribution in whole tissue sections, the whole-slice/section images were analyzed using ImageJ (National Institutes of Health, imagej.nih.gov/ij/), and Qupath (https://qupath.github.io/). Whole slide VSI images were opened in Qupath, an ROI was created for each analyzed sample. The ROI was then exported as raw pixels in TIFF format to imageJ for further quantification. In imageJ, the image was measured for total area of the section, and then converted into an 8-bit image. Image channels red, and green were separated, and subtracted against each other to remove background fluorescence, and non-specific binding, as reported^51,52^. For spinal sections, the threshold was set to an auto thresholder triangle. A macro created in imagej was used to process the data sets. The mean intensity value was divided by the area of the section, and then multiplied by 100 to obtain an intensity percentage of arbitrary units (AU). This was repeated for each section N = 3-5 per brain. The values of each brain section were averaged to obtain a whole brain AU value. The Olympus VS120 Virtual Slide Microscopy was used to identify colocalization of immunolabeling. For colocalization analysis, the VSI image files were opened in Qupath and an ROI was drawn around biologically interesting regions, and sent to imageJ for further analysis. The image channels were separated, and only the red (555 nm), and far red (647 nm) channels were used for colocalization quantification. Each channel had a rolling ball algorithm background subtraction set at 50. Next, the area outside the ROI was cleared. After background was cleared, Coloc 2 (https://imagej.net/plugins/coloc-2) was used to test for Pearson’s correlation, r, and Mander’s coefficient auto threshold, tM1 and tM2. Costes P-Value test was used to verify there was colocalization, and Pearson’s r values was used for final data presentation.

### Kinetics analysis

For each experiment, image sequences were imported into ImageJ. The ROIs, identified from pilot experiments, were the nasal/olfactory bulbs region, spine, sternum, liver, and femur, and the size of each ROI was the same. Mean pixel intensity within each ROI was measured for each time point using the images without enhancement. The background signal was from the same ROI before the injection, and this was subtracted from the intensity at each time point. Maximum intensity was identified as the average intensity over 12 sec at the maximum identified using GraphPad Prism software. The area under the curve (AUC) was determined by running AUC analysis option in GraphPad Prism. The initial linear part of the curve from the intensity profile graph was used to obtain the slope by using linear regression function in the GraphPad Prism.

### Tau-anti-tau complex

The conjugated tau protein was incubated with twice the molar concentration of the antibody (T12) overnight in the fridge, and the same tau concentration was injected.

### Blood samples

At the end of the experiment, blood samples were collected by cardiac puncture (100 μL), and plasma separated by centrifugation. A fixed volume (5 μL) of plasma was used to determine NIRF intensities and for comparison. Background was obtained by using saline at the same volume as used for plasma and the value was subtracted from the plasma values.

### Statistical analysis

Data were analyzed by analysis of variance (ANOVA) followed by post hoc Tukey’s test. Student t-test with Welch’s correction was used for two samples comparison. An outlier test (ROUT method) with Q= 10% was used to remove outliers.

Differences were significant at p < 0.05. For statistical representation, *P < 0.05, **P < 0.01, ***P < 0.001, and ****P < 0.0001 are the levels of statistical significance. NS is not significant. All values were expressed as mean ± SEM. GraphPad Prism software (version 9.5.1 (528)) was used for all analysis.

## Acknowledgements

This work was supported by NIH grants to RD (AG057574) and to the Center for Musculoskeletal Research Histocore (CMSR) core facility (NIH P30 NIAMS AR069655).

## Supplementary Figures

**Fig. S1**. **Blood-derived tau was present in the heart**. Representative image showing that tau-NIRF was associated with the heart 150 mins after its IV injection.

**Fig.S2. H-tau present in bone in Tg-htau-GFP mice. a-b**) Representative images showing htau-GFP in brain and bone of a sagittal head section (**a**), femur and femoral head (**b**), and in liver (**c**). Scale bar = 1000 μm for whole femur section and the sagittal head section, and 200 μm for the other panels.

**Contributions:** AS performed experiments, labeling, imaging, image and data analyses, preparation of figures and videos, contribute to the manuscript (MS) preparation and submission and review the MS. MB performed experiments, imaging, data analysis and prepare figures. AJ, BB and HL performed experiments. NB prepared, purified, and quantified the tau protein samples. NP and CJ contribute to Tg Htau data, RW contributed NIRF resources and reviewed the MS. RK contribute to resources, experimental design and review the MS. RD designed the study and wrote the MS.

**Data sharing**: On request to the corresponding authors, and from Figshares, after publication.

