## supplemental Figures for "Central and peripheral tau retention modulated by an anti-tau antibody"

### Supplementary Figures

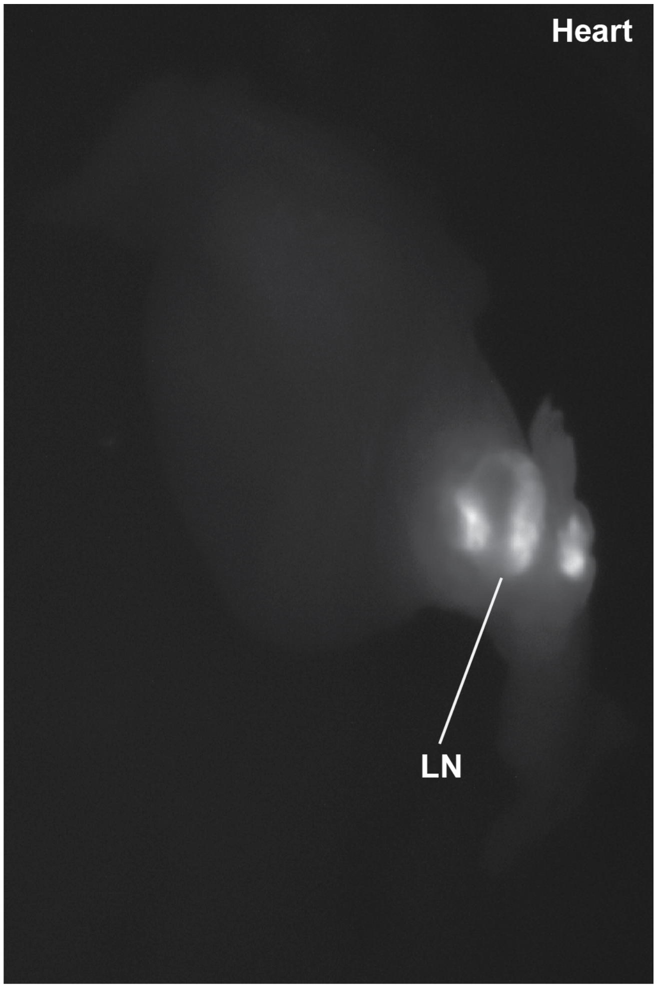

**Fig. S1**. **Blood-derived tau was present in the heart**. Representative image showing that tau-NIRF was associated with the heart 150 mins after its IV injection.

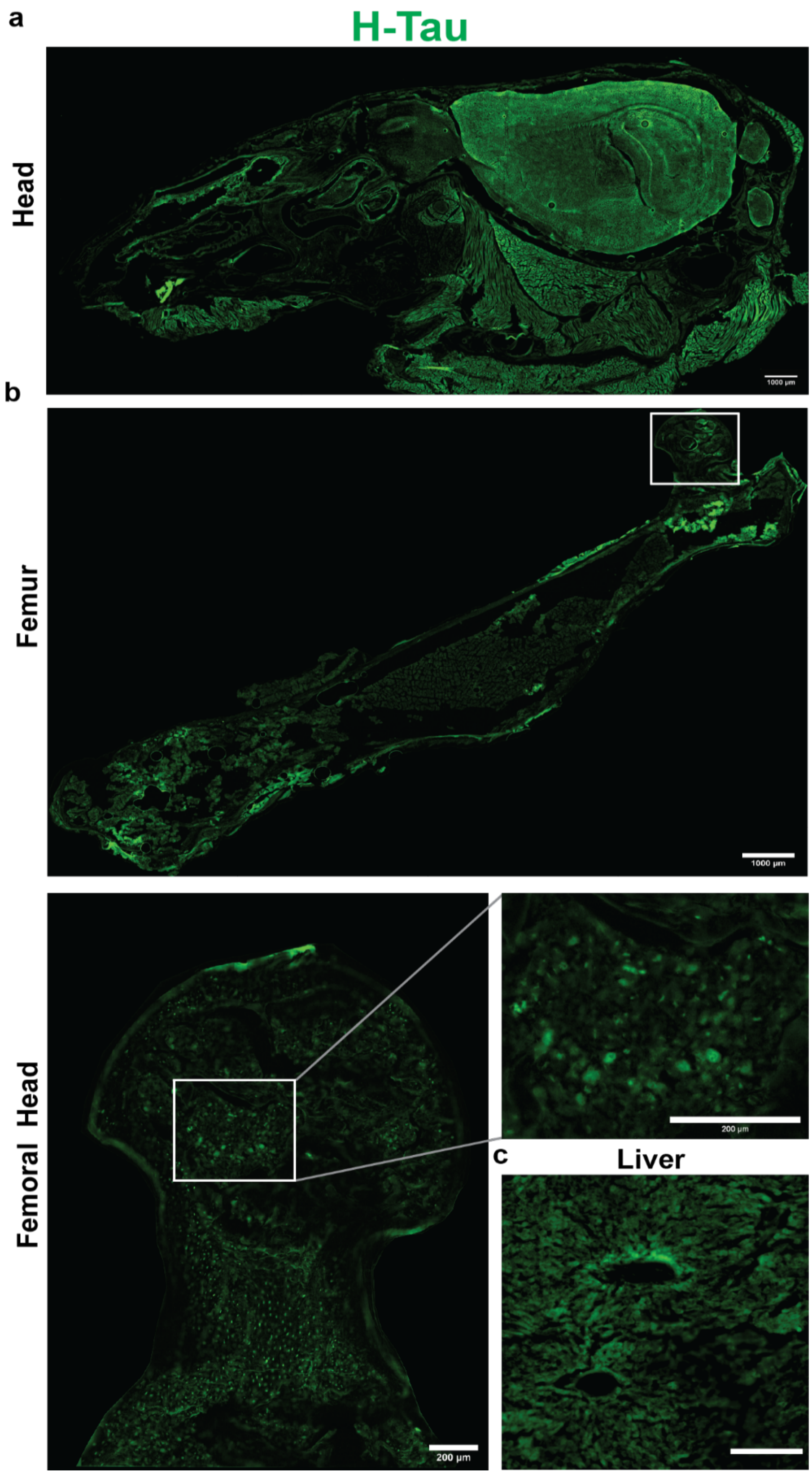
a
H-Tau
Head
1000 µm
b
Femur
1000 µm
Femoral Head
200 µm
200 µm
c
Liver

**Fig.S2. H-tau present in bone in Tg-htau-GFP mice. a-b**) Representative images showing htau-GFP in brain and bone of a sagittal head section (**a**), femur and femoral head (**b**), and in liver (**c**). Scale bar = 1000 μm for whole femur section and the sagittal head section, and 200 μm for the other panels.
